# Zwitterionic hydrogels modulate the foreign body response in a modulus-dependent manner

**DOI:** 10.1101/195719

**Authors:** LE Jansen, LD Amer, E Y-T Chen, TV Nguyen, LS Saleh, TS Emrick, WF Liu, SJ Bryant, SR Peyton

## Abstract

Reducing the foreign body response (FBR) to implanted biomaterials will enhance their *in vivo* performance in tissue engineering. Poly(ethylene glycol) (PEG) hydrogels are increasingly popular for this application due to their low cost and ease of use. PEG hydrogels can elicit chronic inflammation upon implantation, but recent evidence has suggested that extremely hydrophilic, zwitterionic hydrogels can reduce the FBR to particles and gels. To expand on this approach, we synthesized hydrogels of co-monomers PEG and the zwitterion phosphorylcholine (PC) to quantify the combinatorial effects of modulus and hydrophilicity on the FBR. Surprisingly, hydrogels with the highest amount of zwitterionic co-monomer elicited the highest FBR we observed. Lowering the hydrogel modulus (165 kPa to 3 kPa), or PC content (20 wt% to 0 wt%), mitigated this effect. A high density of macrophages was found at the surface of implants associated with a high FBR, and mass spectrometry analysis of the proteins adsorbed to these gels implicated extracellular matrix, immune response, and cell adhesion protein categories as drivers of macrophage recruitment to these hydrogels. Overall, we show that modulus regulates macrophage adhesion to zwitterionic-PEG hydrogels, and demonstrate that chemical modifications to hydrogels should be studied in parallel with their physical properties to optimize implant design.

**Highlights:**

- Modulus and zwitterion content independently modulate the foreign body response to soft hydrogels
- Soft PEG hydrogels synthesized with the zwitterionic PC co-monomer are pro-inflammatory as modulus is increased
- The chemical and physical properties of hydrogels influence the foreign body response via macrophage recruitment and protein adsorption

## 1. Introduction

Poly(ethylene glycol) (PEG) hydrogels are used in tissue engineering because they are compatible with cells and they are easy to chemically functionalize. These features make them attractive biomaterials to control tissue regeneration via porosity [1, 2], stiffness [3, 4], and presentation of peptides and proteins to direct cell growth and migration [5-7]. Though PEG hydrogels have been used extensively *in vitro* to culture chondrocytes [8], mesenchymal stem cells [1], hepatocytes[9], and muscle cells[3], the capacity for these cells to regenerate tissue in PEG could be limited *in vivo* because PEG hydrogels elicit a foreign body response (FBR) [10]. The FBR to implanted materials starts with protein adsorption to the biomaterial, which facilitates macrophage adhesion. Pro-inflammatory macrophages and foreign body giant cells then recruit other cells, such as fibroblasts, to infiltrate and deposit collagen around the implants [11]. This matrix remodeling around hydrogel implants ultimately leads to fibrosis and chronic inflammation. The FBR to PEG hydrogels is proposed to be driven by either its susceptibility to degradation by macrophages [10] or because certain inflammatory proteins can adhere to the hydrogel surface [12]. Since PEG degradation leads to surface fouling and protein adsorption [13], these properties are potentially coupled.

Recent work has shown that certain material properties, like size, shape, stiffness, and charge have a profound impact on the FBR [14-16]. For example, large, spherical hydrogels in a colloidal implant exhibit less fibrosis than small ones [16]. Also, increasing hydrogel stiffness increases the collagen capsule thickness around implants, likely because macrophages are better able to adhere and spread on stiffer hydrogels [17]. Stiffness-driven FBR is a concern, because the ability to regenerate different tissues, like muscle, can rely on stiffness [3]. The FBR could be improved by chemically modifying PEG hydrogels at different stiffnesses. Researchers have shown that including cell adhesive [12] and enzyme degradable peptides [5, 6] reduce the FBR to PEG hydrogels, because they stimulate interactions with the immune system that promote wound healing.

An approach that has been used for other hydrogels, but not PEG, is surface chemical modification to help avoid activating the immune system. For example, a combinatorial approach screened 774 different alginate analogs to find modifications that reduced the FBR to implanted alginate hydrogels by evading recognition [18]. Hydrogels made with the zwitterion carboxybetaine significantly reduced immune response and implant FBR [14]. This study suggests that zwitterionic materials could be an effective way to modulate the FBR. Phosphorylcholine (PC) zwitterions are of particular interest because of their *in vivo* use as medical device coatings [19]. Recently, we synthesized hydrogels composed of PEG and PC, which have an increased range of stiffnesses compared to PEG hydrogels alone, and reduced protein adsorption to the surface *in vitro* [20]. These PEG-PC hydrogels also allow for cell culture on biomaterials of varying stiffness and biochemical complexity [21-23]. We hypothesized that these materials would have a reduced inflammatory response because of minimal protein adsorption, making them potentially advantageous for long-term *in vivo* implants.

To test that hypothesis, we developed a panel of PEG-PC hydrogels with varying stiffness and zwitterion content to investigate how these properties independently contribute to the FBR. Using both *in vitro* and *in vivo* assays, we explored how the chemical and physical compositions of hydrogels influence protein adsorption and macrophage polarization, and ultimately how these properties modulate the FBR.

## 2. Materials and Methods

### 2.1 Hydrogel fabrication

Hydrogels were polymerized as previously described [20]. In brief, PEG dimethacrylate and 2-methacryloyloxyethyl phosphorylcholine (PC) (Sigma-Aldrich, St. Louis, MO) were mixed in PBS at the concentrations shown in Table 1. Pre-polymer solutions were degassed with nitrogen for 30 seconds and cured under UV light (365nm) with 0.8 wt% of Irgacure 2959 (BASF, Ludwigshafen, Germany). For *in vivo* implantation, hydrogels were formed under sterile conditions in a 5 mm diameter by 0.8 mm height cylindrical mold and swollen overnight in PBS (final surface area ranged between 60-120 mm^2^). For *in vitro* studies, hydrogels were swollen in PBS overnight and punched into 6 mm diameter discs with an average height of 0.5 mm.

**Table 1.** Polymer composition and Young’s modulus for hydrogel implants

| Gel Name | PC (wt%) | PEG (wt%) | Modulus (kPa)<br>± SD |
| --- | --- | --- | --- |
| A | 20 | 1 | $3 \pm 1$ |
| B | 20 | 4 | $64 \pm 4$ |
| C | 20 | 6 | $165 \pm 22$ |
| D | 15 | 8 | $169 \pm 25$ |
| E | 8 | 12 | $161 \pm 45$ |
| F | 0 | 18.4 | $174 \pm 34$ |

### 2.2 In vivo hydrogel implants

Hydrogel disks were implanted subcutaneously into the dorsal pockets of eight-week-old C57BL/6 male mice (Charles River Laboratories, Wilmington, MA) by making a subcutaneous incision along the centerline of the back shoulder blades. Hydrogels were left *in vivo* for 30 minutes for short-term protein adsorption studies and for 28 days for quantification fibrous capsule formation and macrophage recruitment. Each animal received four implants, each implant consisting of unique hydrogel chemistry. Both hydrogel chemistry and location of biological replicates were randomized on the backs of each mouse (N=4 for each variation of hydrogel implanted). Endotoxin levels were measured using the ToxinSensor Chromogenic LAL Endotoxin Assay Kit in accordance with the manufacturer’s instructions (Genscript, China).

### 2.3 Animal protocols

All animal protocols were in accordance with NIH guidelines for animal handling and approved by the University of California Irvine, the University of Massachusetts Amherst, and the University of Colorado at Boulder Institutional Animal Care and Use Committees.

### 2.4 Protein adsorption

Hydrogel implants were explanted after 30 minutes of incubation *in vivo* to determine initial protein adsorption to gel surfaces. Hydrogels were incubated for 30 minutes in 10% fetal bovine serum (FBS) (Thermo Fisher Scientific, Waltham, MA) in PBS for *in vitro* protein adsorption. To remove and quantify the adsorbed proteins, hydrogels were soaked in 50 mM ammonium bicarbonate (Thermo) solution for 2 hours. Solutions were removed and either immediately processed or flash frozen and stored in the -80°C until processing. *In vitro* gels were additionally incubated with 1 wt% SDS (Hoefer, Holliston, MA) for 30 minutes, and solutions were immediately processed. Total protein concentration was measured using a bicinchoninic acid assay (BCA) assay according to manufacturer’s instructions (Thermo). Protein was loaded at 10 μg/lane, run on a 4-29% tris-glycine polyacrylamide gel, stained using silver stain according to manufacturer’s instructions (Thermo), and imaged using the IN Genius Syngene Bioimaging platform (Frederick, MD).

### 2.5 Mass Spectrometry

Proteins removed from the explanted hydrogels were reduced in 10 mM dithiothreitol (DTT) (Thermo) for 30 minutes at 37°C. Samples were alkylated with 20 mM iodoacetamide (Sigma-Aldrich) in the dark at room temperature for 30 minutes. The solution was quenched with 5 mM DTT prior to cleavage. Proteins were cleaved via trypsin (Thermo) and Lys-C endoproteinase (Promega, Madison, WI) at a ratio of 1:50 enzyme to protein overnight (12-16 hours) at 37°C. A reverse phase LC gradient was used to separate peptides prior to analysis. Mass spectrometry analysis was performed in an Orbitrap Fusion Tribrid (Thermo). Peptides were aligned against the UniProt *Mus musculus* proteome using the Thermo Proteome Discoverer 1.41.14. Parameters used trypsin as a protease, with 4 missed cleavages per peptide, a precursor mass tolerance of 10 ppm, and fragment tolerance of 0.6 Da. Proteins identified had a PSM>1, coverage >10%, unique peptides >1, and a protein score >0 (protein score is developed by Proteome Discoverer to indicate confidence for a protein hit). Of these hits, only full-length proteins identified on at least two of the three hydrogel replicates are reported for each sample condition. Hierarchical clustering analysis was performed using the MATLAB Bioinformatics toolbox R2015b (Mathworks, Natick, MA) on the proteins removed from explanted hydrogels and identified through mass spectrometry. Euclidean distance and average linkage were used to generate the dendrogram. Data were normalized by giving proteins present on the hydrogel a value of 1 and proteins absent a value of 0.

### 2.6 Gene Ontology

The Database for Annotation, Visualization, and Integrated Discovery (DAVID) v6.7 (http://david.abcc.ncifcrf.gov/) [24, 25] was used to assess the biological process and cellular component gene ontology terms associated with each of the identified proteins. All proteins identified across the substrates were submitted as background and the individual protein hits for each hydrogel were compared to find gene ontology groups. Notable ontology groups are highlighted, and the p-values are reported.

### 2.7 Histological analysis

Swartzlander et al. have previously described the tissue preparation, imaging, and image analysis used here in detail [12]. Briefly, hydrogels were explanted alongside the dorsal skin, fixed in paraformaldehyde, and embedded in paraffin. Samples (10 μm thick) were stained with Masson’s Trichrome via standard protocols. Collagen density was measured using the protocol published by Zhang et al. [14]. Macrophages were stained with the rat anti-mouse Mac3 as the primary antibody (1:30, Abcam, Cambridge, MA) and the biotin anti-rat IgG (1:30, Abcam).

### 2.8 Primary macrophage adhesion

Monocytes were isolated from the bone marrow of 7 to 10 week-old C57BL/6 male mice (Jackson Laboratory, Bar Harbor, ME) as described previously [26]. Cells were separated using Lympholyte M (Accurate Chemical, Westbury, NY) and plated in macrophage differentiation medium (IMDM with 20% FBS, 2 mM L-glutamine (Thermo), 1% penicillin-streptomycin (Corning, Corning, NY), 1.5 ng/ml recombinant mouse macrophage colony stimulating factor (M-CSF, R&D Systems, Minneapolis, MN), and 100 ng/ml flt-3 ligand (R&D Systems) for 5 days. Macrophages were lifted from culture plates using a cell scraper and 0.05% trypsin-EDTA. A soybean trypsin inhibitor (Thermo) was used in place of serum. Cells were seeded at 30,000 cells/cm^2^ in serum-free medium on hydrogels swollen in PBS or in 10% FBS in PBS, Human Plasma Fibronectin (Millipore, Billerica, MA), Collagen I from rat tail (Thermo), or active mouse Fibrinogen protein (ab92791, Abcam) for 30 minutes prior to seeding. After 24 hours, hydrogels were rinsed, fixed in 4% formaldehyde, and adhered macrophages were stained with DAPI at 1:10000 (Thermo). Adhered macrophages were imaged on a Zeiss Axio Observer Z1 microscope (Carl Zeiss, Oberkochen, Germany) using an AxioCam MRm camera and an EC Plan-Neofluar 20X 0.4 NA air objective and manually quantified using ImageJ (NIH, Bethesda, MD).

### 2.9 Assessment of cytokine secretion

Bone marrow-derived macrophages used in the cytokine secretion assay were harvested from the femurs and tibias of 6–10 week-old C57BL/6 mice (Jackson Laboratory) as previously described [27]. Briefly, cells were treated with ACK lysis buffer (Thermo), centrifuged, and resuspended in culture medium containing 10% heat-inactivated FBS and recombinant M-CSF for macrophage differentiation. BMDM were dissociated using cell dissociation buffer (Invitrogen) and scrapers on day 6-8 and seeded on the gels in culture media. At hour 6 after cell seeding, the culture media was replaced with one of the four conditioning media: 1) no stimulation, 2) 1 ng/mL of LPS/IFN-D (Sigma and BioLegend, San Diego, CA) each, 3) 20 ng/mL IL-4/IL-13 (BioLegend) each, and 4) 0.5 ng/mL LPS and 20 ng/mL IL-4/IL-13 each. After 12 hours of incubation, the supernatants were collected and analyzed for TNF-α and IL-10 secretion by enzyme-linked immunosorbent assay (ELISA) following the manufacturer’s instructions (BioLegend).

### 2.10 Statistical Analysis

Statistical analysis was performed using GraphPad’s Prism v7.0a (La Joya, CA). Statistical significance was evaluated using either a two-tailed t-test was used or a one-way analysis of variance where noted, followed by a Tukey’s post-test for pairwise comparisons. Spearman correlations were calculated from means and standard deviations paired by the condition. P-values <0.05 are considered significant, where p<0.05 is denoted with *, ≤0.01 with **, and ≤0.001 with ***.

## 3. Results

### 3.1 The zwitterion phosphorylcholine increases the FBR to PEG hydrogels

We explored the interplay between hydrogel modulus and zwitterionic content, two parameters that separately regulate immune response to hydrogel implants. Using our PEG-PC hydrogel [20], we created a panel of conditions that independently modulated either hydrogel modulus or zwitterion content (Table 1). Hydrogel stiffness, measured using bulk compression rheology, was increased from ∼3 to 165 kPa while PC content was kept at 20 wt% or held constant at 165 kPa while PC was decreased from 20 to 0 wt% (Figure 1a-b). The soft, high PC hydrogels swelled the most, and hydrogel swelling decreased with increasing stiffness and removal of the PC zwitterion (Figure 1c). These hydrogels, which had low endotoxin levels (<0.08 EU/mL), were implanted into C57BL/6 mice for 8 weeks. We first observed that with a fixed zwitterion content, increasing the stiffness increased the thickness of the fibrous capsule, a result consistent with previous findings (conditions A-C in Figure 1d-e) [17]. Surprisingly, reducing the amount of zwitterion, while holding the modulus of the hydrogels at the highest value we tested, decreased the fibrous capsule thickness (conditions C-F in Figure 1d-e). Though both hydrogel stiffness and PC content contributed to the final thickness of the collagen capsule, the collagen density throughout the capsule was not significantly different among any of the hydrogels (Suppl. Figure 1).

**Figure 1.**
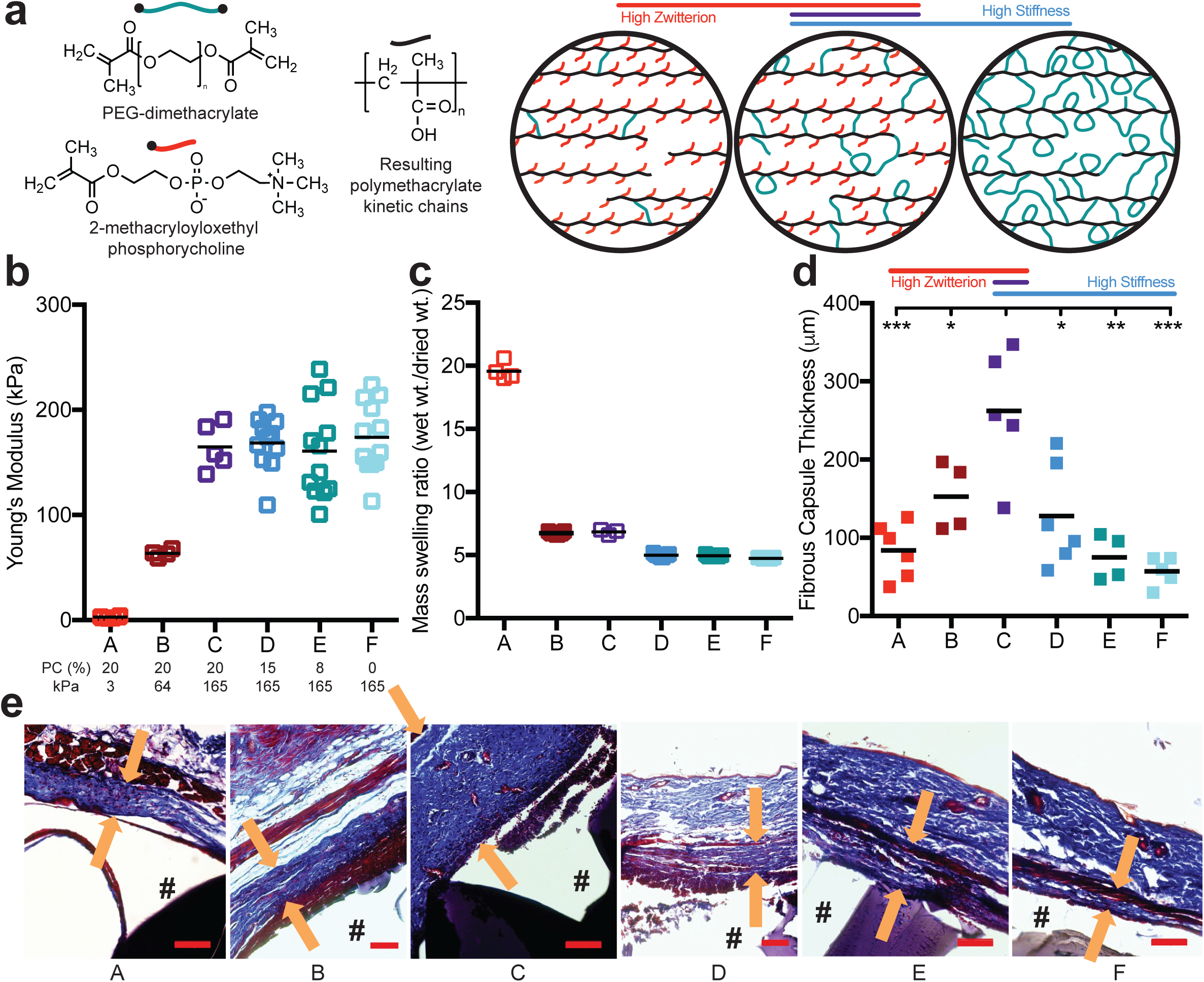
The FBR to PEG hydrogels is highest on stiff, highly zwitterionic implants. a) A schematic of the hydrogel composed of PEG-dimethacrylate (PEG, green) and 2-methacryloxyloxethyl phosphorycholine (PC, red) to produce polymethacrylate chains (black) with pendant PC groups and PEG crosslinks. b) The Young’s modulus and c) mass swelling ratio of the different hydrogel conditions as both PEG and PC content was modulated (N≥4). The molar percentage of PC and the Young’s modulus for each hydrogel is labeled below b. d) 28 days after hydrogels were subcutaneously implanted into a mouse, the fibrous capsule thickness was measured using a Masson’s Trichrome stain (N≥4). e) Representative images for each stain, where “#” denotes the location of the hydrogel, and the arrows represent the thickness measurement taken (scale bar = 100 μm). Significance was determined using an ANOVA with Tukey’s post-test, where p=0.05 was considered significant.

### 3.2 Proteins associated with the extracellular matrix are enriched on the high FBR hydrogel

One of the first steps in the FBR is the formation of a provisional protein matrix[11]. We previously showed that the addition of PC to PEG hydrogels decreases protein adsorption to hydrogels *in vitro* [20], and decreasing non-specific protein adsorption with zwitterions has been proposed by others to reduce the FBR [14]. However, because the PEG-PC hydrogels we synthesized have a maximum amount of 20 wt% zwitterion, non-specific protein adsorption likely still occurs. We quantified the total protein adsorbed to the surface of our hydrogels (Table 1) during *in vitro* exposure to serum to see if this explained our observation of the enhanced FBR on the stiffest, highly zwitterionic hydrogels. In both *in vivo* and *in vitro* experiments, we observed that the softest hydrogel condition, which exhibited the lowest FBR, had the most total protein accumulated (Figure 2a-b). Thus, total adsorbed protein could not explain the FBR to our array of hydrogels.

**Figure 2.**
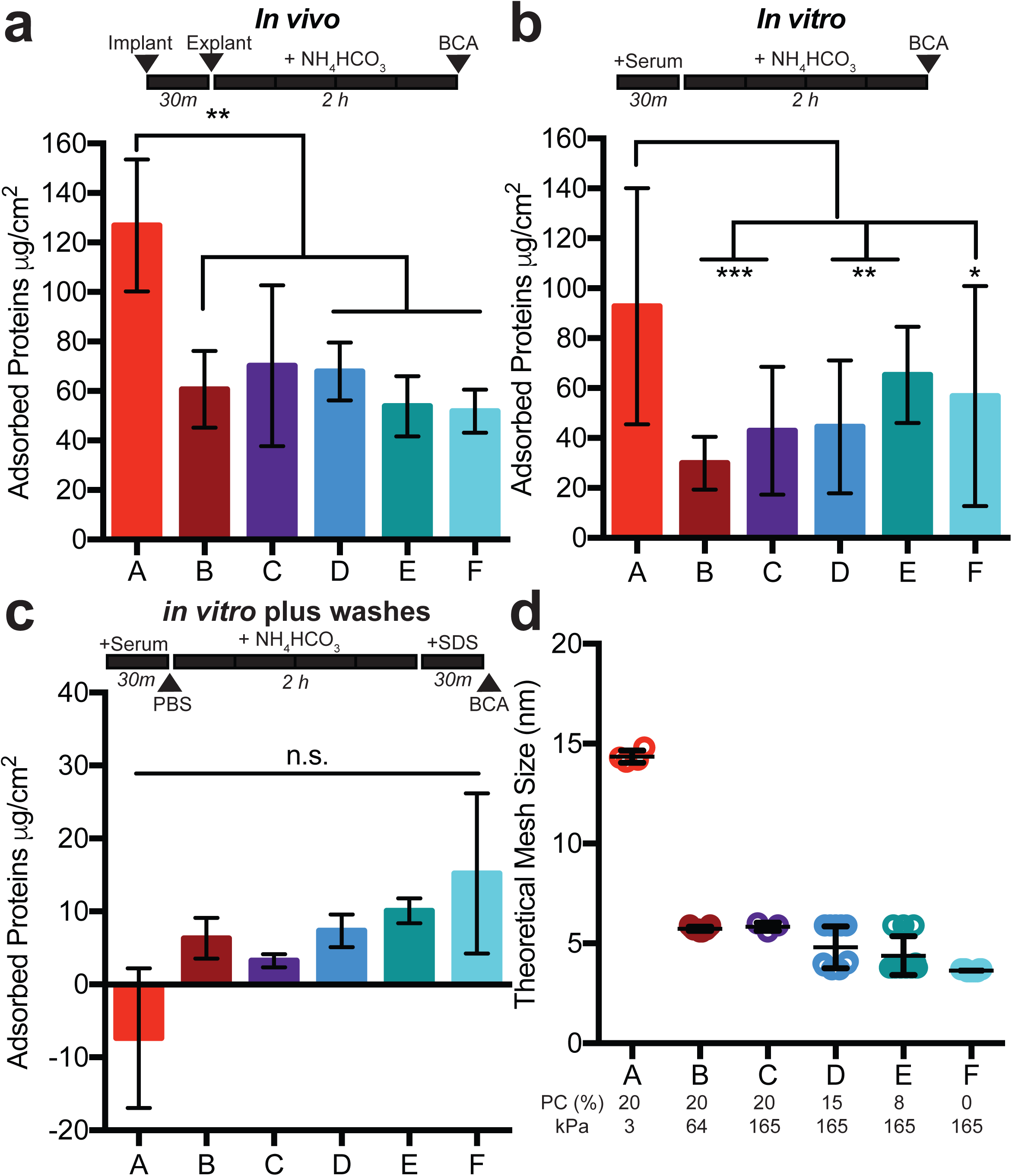
Total protein adsorption to hydrogels is not sufficient to explain the FBR. A protein assay was used to quantify the amount of protein adsorbed to each hydrogel after a) *in vivo* implantation, b) exposure to 10% serum protein in PBS *in vitro*, or c) with a PBS and 1wt% SDS wash added after exposure to 10% serum proteins *in vitro*. All hydrogels were exposed to ammonium bicarbonate (NH_4_HCO_3_) for 2 hours before analysis with a bicinchoninic acid assay (BCA). The wash timeline is depicted above each graph. d) The theoretical mesh size of each hydrogel calculated using the Flory theory as modified by Canal and Peppas. The percentage of PC content and the Young’s modulus for each hydrogel is labeled below. Significance is determined using an ANOVA with Tukey’s post-test where p=0.05 is significant. Error bars are the SD (N=2, n=3).

Both of these methods demonstrated that this zwitterionic content did not decrease the total amount of protein adsorbed to hydrogels, which contradicted our original hypothesis. We then used a series of more rigorous rinse methods to determine if the degree of protein binding to different surfaces was confounding our results. When a phosphate buffered saline (PBS) wash was added before protein removal with ammonium bicarbonate (Suppl. Figure 2c), we found that ∼75% of the total protein was removed, indicating that most of the protein adsorbed passively on the hydrogel surface and was not tightly bound. We also added a final wash with sodium dodecyl sulphate (SDS), and found that proteins, though minimal, could be detected on the hydrogels post-washing with ammonium bicarbonate, and that the addition of PC reduced how strongly the proteins adsorbed to the hydrogels (Figure 2c). We then speculated that smaller proteins might be diffusing into the softest, most porous hydrogels (condition A in particular, see Figure 1c and 2d). In fact, the mesh size of our hydrogels correlated with the total protein adsorption during gentle rinsing (Figure 2d, Spearman ρ=0.94, p=0.016), suggesting that larger pore sizes trapped small proteins within the mesh. Overall, these results indicate that the increased protein adsorption (Figures 2a-b) is a larger indication of small proteins diffusing into the network, rather than more proteins adsorbing to the gel surfaces (Figure 2c), and that zwitterions likely impact how tightly bound proteins are to the surface.

Since total protein adsorption did not correlate with the observed FBR, we next examined whether the identity of the proteins adhered to the hydrogels could predict the FBR. Using a silver stain, we were unable to distinguish any major differences in the protein molecular weight signature adsorbed to the implanted *in vivo* hydrogels (Suppl. Figure 3a). From the more sensitive liquid chromatography-mass spectrometry (LC-MS) technique, we annotated over 350 of the proteins that adsorbed to each hydrogel surface 30-minutes after *in vivo* implantation (Figure 3a, Suppl. Table 1). The majority of the top 20 protein hits were conserved across all the hydrogels, in agreement with our silver stain results (Suppl. Table 2, Suppl. Figure 3b). Unsupervised hierarchical clustering of protein hits identified using LC-MS separated hydrogels by either high stiffness or high zwitterion content (Figure 3b, separation indicated by Euclidean distance). LC-MS was also unable to identify any appreciable trends in protein molecular weights adhered to each hydrogel (Suppl. Figure 3c). However, a higher percentage of proteins with extreme isoelectric points (above 9 and below 5) adsorbed to the stiffer hydrogels containing PC, compared to hydrogels with low or no PC included (Suppl. Figure 3d-e).

**Figure 3.**
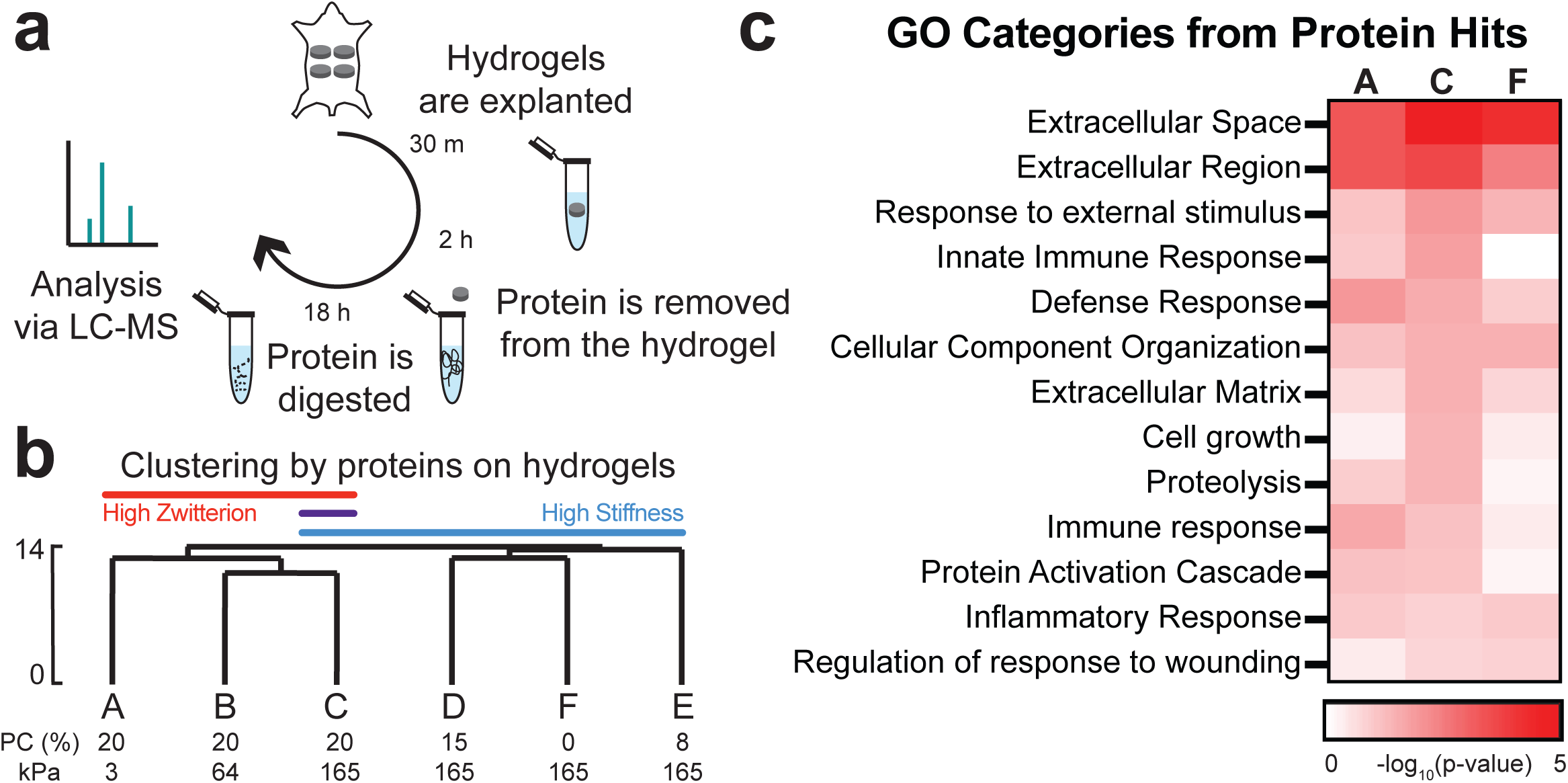
Identity of adsorbed proteins distinguishes highly zwitterionic from stiff hydrogels. a) Schematic of how protein was collected from implants and analyzed via LC-MS. b) Hierarchical clustering of LC-MS data normalized by the type of protein adsorbed to each hydrogel. The percentage of PC content and the Young’s modulus for each hydrogel is labeled below. The scale depicted on the side is the Euclidean distance. c) Heat map of the log_10_(p-value) for select Gene Ontology (GO) categories identified from the proteins that adhered to hydrogels A, C, and F. These were identified using DAVID with all proteins identified as the background.

It is well known that hydrogel surface chemistries can influence protein adsorption [28, 29]. Given that clustering of proteins distinguished the higher zwitterionic from the stiffest hydrogel conditions, we hypothesized that a subset of proteins could be identified that were recognized by immune cells and drove the observed FBR. A DAVID analysis identified categories of proteins associated with biological processes and cell components that adsorbed onto the hydrogels with the maximum (C) and minimum (A and F) collagen capsule thickness. We focused our search on categories associated with cell and immune response, extracellular matrix (ECM), and ECM remodeling because we speculated these categories would help to explain the formation of the fibrous capsule. In fact, a higher percentage of proteins from these biological categories were found on the high FBR hydrogel compared to the others, as confirmed by more significant p-values (Figure 3c). This suggests that these adsorbed proteins may be driving macrophage recruitment and promoting ECM remodeling on the surface of the stiffer, more zwitterionic hydrogels. This might in part explain be why this particular material exhibited heightened inflammation *in vivo*.

### 3.3 Stiffer hydrogels with higher PC promote an inflammatory phenotype in macrophages

Macrophages are one of the most prominent cell types that accumulate rapidly to the surface of implanted materials and devices, and they play a major role in initiating chronic inflammation [30]. We hypothesized that macrophage adhesion to the proteins we detected via LC-MS may drive the observed FBR response to the stiffer, more zwitterionic hydrogels. Therefore, we examined macrophages present near the implants *in vivo* and performed a macrophage adhesion experiment *in vitro.* The extent of macrophage infiltration around implanted hydrogels *in vivo* was highest around the high FBR hydrogel (condition C, Figure 4a). Similarly, macrophage adhesion to these hydrogels *in vitro* was highest on this hydrogel condition (Figure 4b). Interestingly, this *in vitro* adhesion observation was only true in the presence of serum, suggesting that protein adsorption to the surface of the gels is required for adhesion and initiating the FBR.

**Figure 4.**
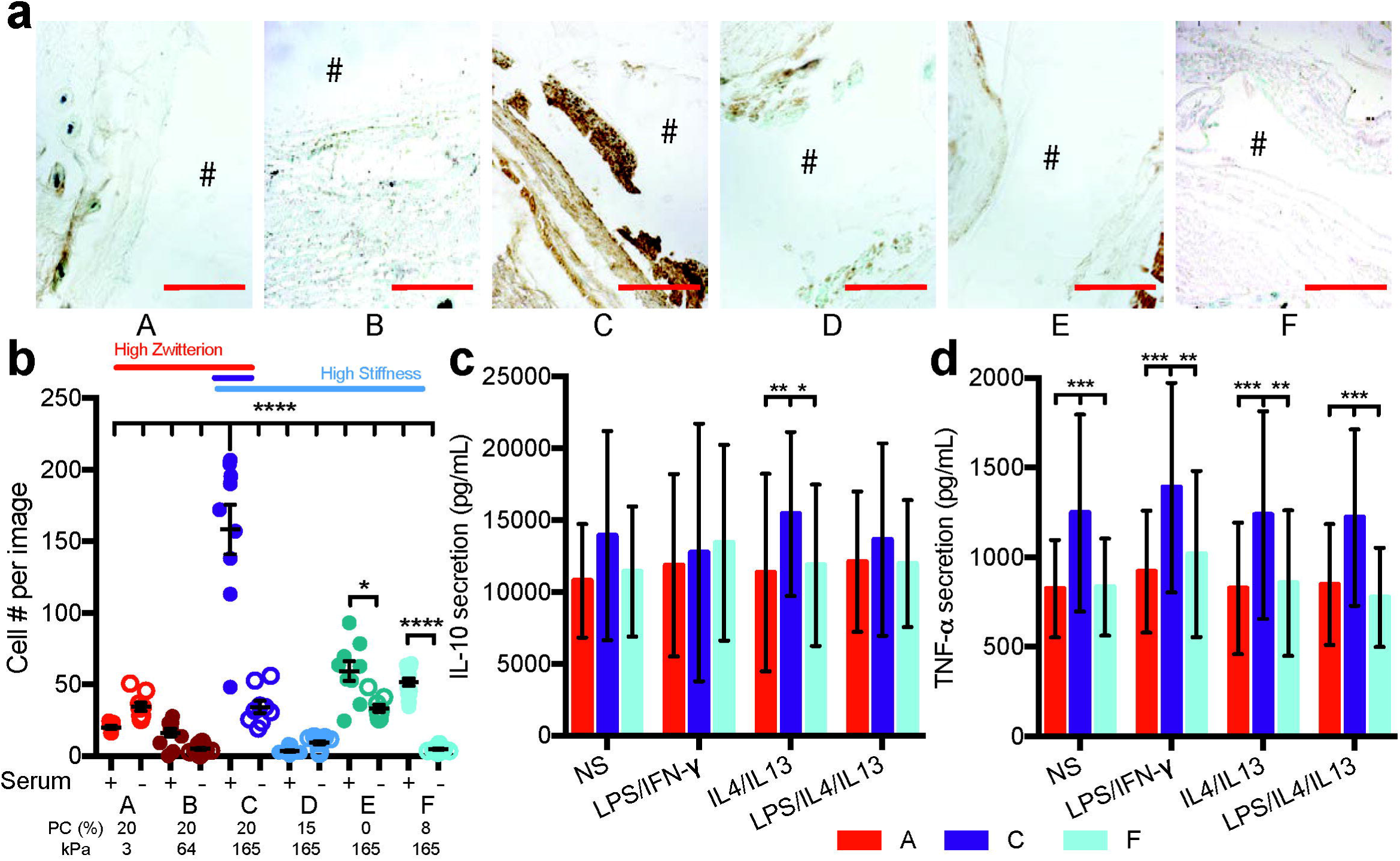
Macrophages adhere better to implants with a more severe foreign body response. a) Representative images for macrophages stained around the implant, where the # denotes the location of the implant (scale 250 μm). b) Cell adhesion to hydrogels either treated for 30 minutes with serum proteins or not and then seeded with macrophages and imaged 24 hours later (N=2, n=4). Stats displayed for plus and minus serum on hydrogels and hydrogel C plus serum compared to all other hydrogels. Secretion of c) IL-10 and d) TNF-alpha from macrophages seeded on hydrogels with different stimulation factors. Stimulation of each condition was as follows: NS: no stimulation, LPS/IFN-γ: 1ng/mL LPS and IFN-γ, IL4/IL13: 20ng/mL IL4 and IL13, LPS/IL4/IL13: 0.5ng/mL LPS and 20ng/mL IL4 and IL13. The percentage of PC content and the Young’s modulus for each hydrogel is labeled below. Significance is determined using an ANOVA with Tukey’s post-test where p=0.05 is significant. Error bars are the SD (N=2,n=5).

During the FBR, adhered macrophages recruit fibroblasts to the wound site to begin ECM turnover using cytokines [11]. Thus, we examined for inflammatory cytokines secreted by macrophages attached to either the high FBR or the two lowest FBR hydrogels *in vitro* (conditions C vs. A and F, Figures 4c-d). Secretion of TNF-α is associated with pro-inflammatory macrophages, and IL-10 is associated with anti-inflammatory macrophages [31]. Macrophages secreted high quantities of IL-10, regardless of hydrogel condition (Figure 4c), but TNF-α secretion was highest on the hydrogel condition that produced the highest FBR (Figure 4d). Interestingly, cytokine secretion was independent of the polarizing stimuli added to the medium including LPS, IFN-γ, IL4, and IL13, which were confirmed to influence the amount and type of cytokine secreted when cells are cultured on tissue culture plastic (Figure 4c-d, Suppl. Figure 4a-b).

## 4. Discussion

Although PEG-based materials have been used widely in tissue engineering applications [32, 33], recent studies have shown they elicit an inflammatory response [10]. This response could be reduced using zwitterions [14], one example being zwitterion-coated nanocarriers [34]. Here, we investigated if the zwitterionic monomer PC could reduce the FBR when incorporated into PEG hydrogels.

We developed a panel of hydrogels with varying amounts of PEG and PC to independently modulate stiffness and zwitterionic content (Figure 1a-c). Reducing the amount of PC in hydrogels with a constant bulk modulus decreased the fibrous capsule thickness, whereas increasing the amount of PEG crosslinker in hydrogels with a constant amount of PC increased the capsule thickness (Figure 1d-e). Stiffness-driven inflammation has been previously reported in PEG hydrogels, where increased crosslinking promoted increased macrophage spreading on the surface [17]. Surprisingly, the higher modulus hydrogels made without zwitterion had a significantly lower FBR compared to the PEG-PC hydrogels. Others have reported that zwitterionic hydrogels reduce FBR [14], but this previous work utilized much stiffer hydrogels than those explored here and did not study the impact of zwitterions when other physical properties of hydrogels were varied. This is an important feature as stiffness is now well appreciated to drive cell behavior and can impact the ease of materials handling by a surgeon. In light of our results, we propose that the stiffness of the material changes the immune response to the zwitterion PC by modulating protein adsorption and macrophage adhesion associated with FBR, and this should be tested for other ranges of stiffness and PC content. Though the higher PC content hydrogels at low stiffness had lowered immune response, incorporating degradable motifs [6], binding ligands [12], or encapsulated cells [35] into PEG hydrogels might be more effective to modulate the FBR on hydrogels with higher moduli.

The thick layer of macrophages surrounding the high FBR hydrogel (Figure 4a) correlated with the collagen capsule thickness around the implant (Figure 1d). We further explored these results *in vitro* and observed that adhesion of macrophages was highest on our high FBR hydrogel (Figure 4b). The adhered macrophages also expressed significantly more TNF-α on this hydrogel compared to the two lowest FBR conditions (Figure 4c), suggesting that the adhered macrophages were more pro-inflammatory. These data demonstrate that hydrogel C promoted the most pro-inflammatory phenotype in macrophages, which may have caused them to orchestrate the highest FBR.

Hydrogel surface properties can influence macrophage phenotypes. For example, adding an RGD integrin-binding motif to PEG promotes macrophage adhesion and wound healing, effectively decreasing the collagen capsule around hydrogel implants [12]. Interestingly, stimulating macrophages with cytokines used to direct cell phenotype did not influence the cell expression of TNF-α when seeded on hydrogels, but did impact cell phenotypes on tissue culture plastic (Figure 4c-d, Suppl. Figure 4a-b). This result indicated that the surface properties of the hydrogels were more important contributors to the macrophage phenotype than the cytokine cocktails typically used in the field. Additionally, because these hydrogels are not fully zwitterionic, we hypothesized that the surface chemistry of the material influenced protein absorption, because cell adhesion increased in the presence of serum (Figure 4b). Increased cell adhesion did not correlate with an increase in the total protein adsorbed on the hydrogel surface (Figure 4b and Figure 2), but instead was likely dictated by the specific subset of proteins adsorbed that we detected via LC-MS (Figure 3). Recent work has identified specific proteins from serum, like clusterin, that make nanocarriers stealthy *in vivo* [36] and others have shown that coating materials with specific proteins can inhibit the inflammatory response of macrophages [27, 37]. Thus, protein type might be more influential to the FBR than total protein amount.

LC-MS performed on proteins stripped off implanted hydrogels identified over 350 proteins adsorbed to these hydrogels *in vivo*. Many of the protein hits were shared across all the hydrogels screened (Suppl. Figure 3a,b, Suppl. Table 2). Our most abundant protein hit was albumin (Suppl. Table 1), known as the major protein that adsorbs to surfaces of implants and/or injected nanoparticles [12, 36, 38]. Hierarchical clustering of all the proteins separated the higher stiffness and higher PC content hydrogels. We speculate that this separation was driven by the repulsion of highly charged proteins on hydrogels with the higher PC content (Suppl. Figure 3d). However, the differences between the types of proteins adhering to the hydrogels was greater than the proteins separating high and low PC hydrogels, indicated by the Euclidean distance. This highlighted that both the stiffness and the charge displayed on the hydrogel surface influence the provisional proteins adsorbed. Unsurprisingly, other hits, like Vitamin D binding protein [39], Apolipoprotein A-I or A-IV [40], and hemopexin [41], are known to be associated with inflammation, which was likely initiated during the implant procedure. The protein clusterin, which has been implicated in helping nanocarriers avoid macrophage uptake, was found on all hydrogel surfaces, and was most abundant on the two lowest FBR conditions A and F (Suppl. Table 2), potentially explaining why macrophages expressed less TNF-α on this surface. Additionally, fibrinogen α, β, and γ were detected on all our hydrogels except the higher stiffness, more zwitterionic hydrogel (condition C) and, interestingly, fibrin matrices promote IL-10 secretion over TNF-α [31].

Gene ontology on the protein hits revealed that categories associated with ECM, immune response, and cell adhesion were most associated with the high FBR conditions (Figure 3c). This suggests that the provisional matrix that assembled on these materials may facilitate the initial adhesion of macrophages and thereby direct the long-term immune response. We independently screened three different ECM proteins, identified in the protein hits, and found that they assisted with macrophage adhesion to our PEG-only hydrogel (Suppl. Figure 4c). Blood plasma does not follow the same fouling principles on PEG as single or binary protein solutions [42], potentially explaining why these single proteins did not produce the same adhesion result as the complex serum. In fact, protein display can influence how macrophages respond. For example, fibrinogen can stimulate macrophage inflammation in a soluble form but inhibit inflammation when displayed as a fibrin matrix [31]. Our data shows that many of our top protein hits are consistent across the hydrogels (Suppl. Table 2), so the difference we see in macrophage response might be how individual proteins are displayed rather than protein identity. Since the mechanism of protein fouling is different on hydrophilic versus hydrophobic surfaces [43, 44], and we studied a range of hydrophilic hydrogels here, we speculate that the mechanism of fouling may have influenced cell polarization via changes in protein display.

## 5. Conclusion

The FBR to PEG-PC hydrogels changes with respect to both the stiffness and zwitterionic content studied here. Our high FBR hydrogel promoted an inflammatory macrophage phenotype *in vitro*, which agreed with the *in vivo* collagen capsule thickness. We speculate that controlling which proteins adsorb to the material surface within the first 30 minutes of implantation is critical in modulating the FBR to PEG-based hydrogels. While many parameters can confound our understanding of the FBR to implanted biomaterials, this work demonstrates that both stiffness and zwitterion content can independently modulate the FBR. Overall, we identified that the physical properties of implanted materials should be studied in conjunction with chemical surface modifications to fully understand the subsequent immune response.

## Acknowledgements

This work was funded by an NIH New Innovator Award 1DP2CA186573-01 awarded to SRP and 1DP2DE023319-01 to WFL, and the NIH award #1R21AR064436 awarded to SB. SRP is a Pew Biomedical Scholar supported by the Pew Charitable Trusts. TVN and SRP were supported by a Barry and Afsaneh Siadat Career Development Award. LSS was supported by a graduate assistantship in areas of need from the Department of Education. Research reported in this publication was supported by the NIH Award #S10OD010645. The content is solely the responsibility of the authors and does not necessarily represent the official views of the National Institutes of Health.

## Author Contributions

All authors contributed to experimental design, data analysis, and wrote the manuscript. LEJ, EY-TC, TVN, LSS, and LDA performed all the experiments.

## Competing financial interests

The authors declare no competing financial interests.

---

**Figure S1.**
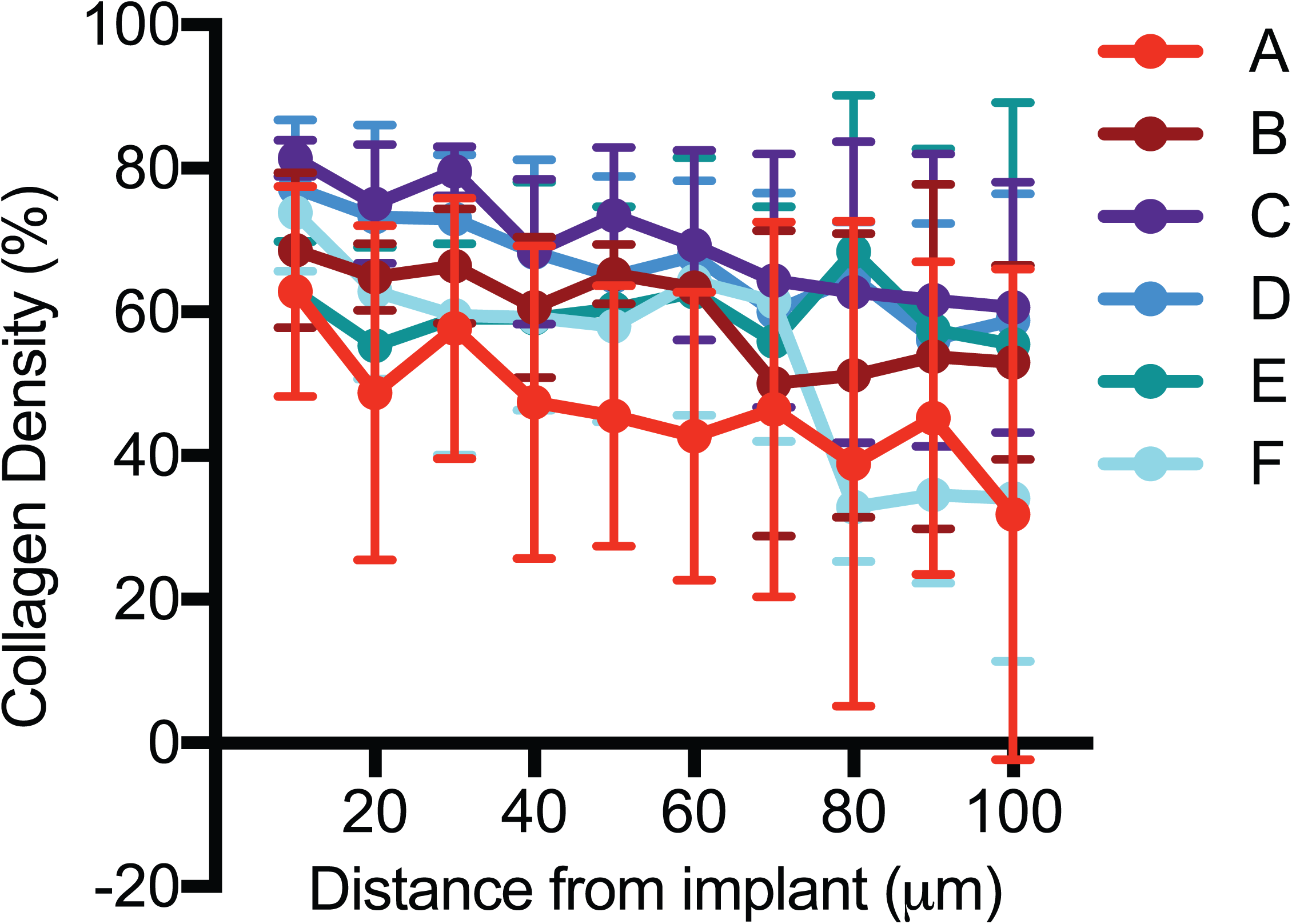
Collagen density was analyzed within 100 μm from the interface of each hydrogel implant using image analysis. Error bars are the SD (N≥4).

**Figure S2.**
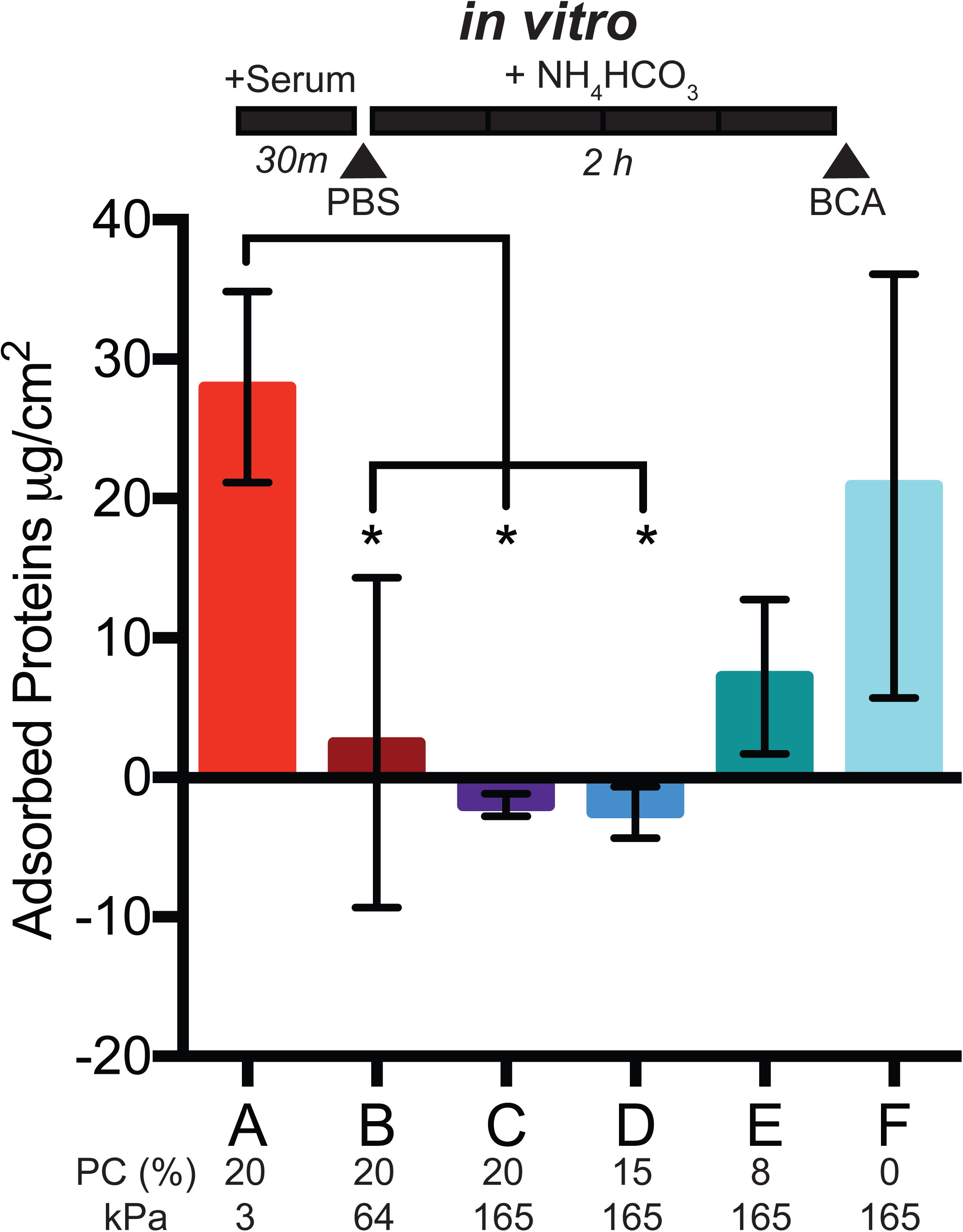
A protein assay was used to measure amount of adsorbed protein onto hydrogels pretreated with 10% serum proteins (FBS) in PBS for 30 minutes. The timeline for the experiment is above the graph. A PBS wash was included before a 2-hour wash with ammonium bicarbonate (NH_4_HCO_3_) and analysis with a bicinchoninic acid assay (BCA). The percentage of PC content and the Young’s modulus for each hydrogel is labeled below. Significance is determined using an ANOVA with Tukey’s post-test where p=0.05 is significant.Error bars are the SD (N=2, n=3).

**Figure S3.**
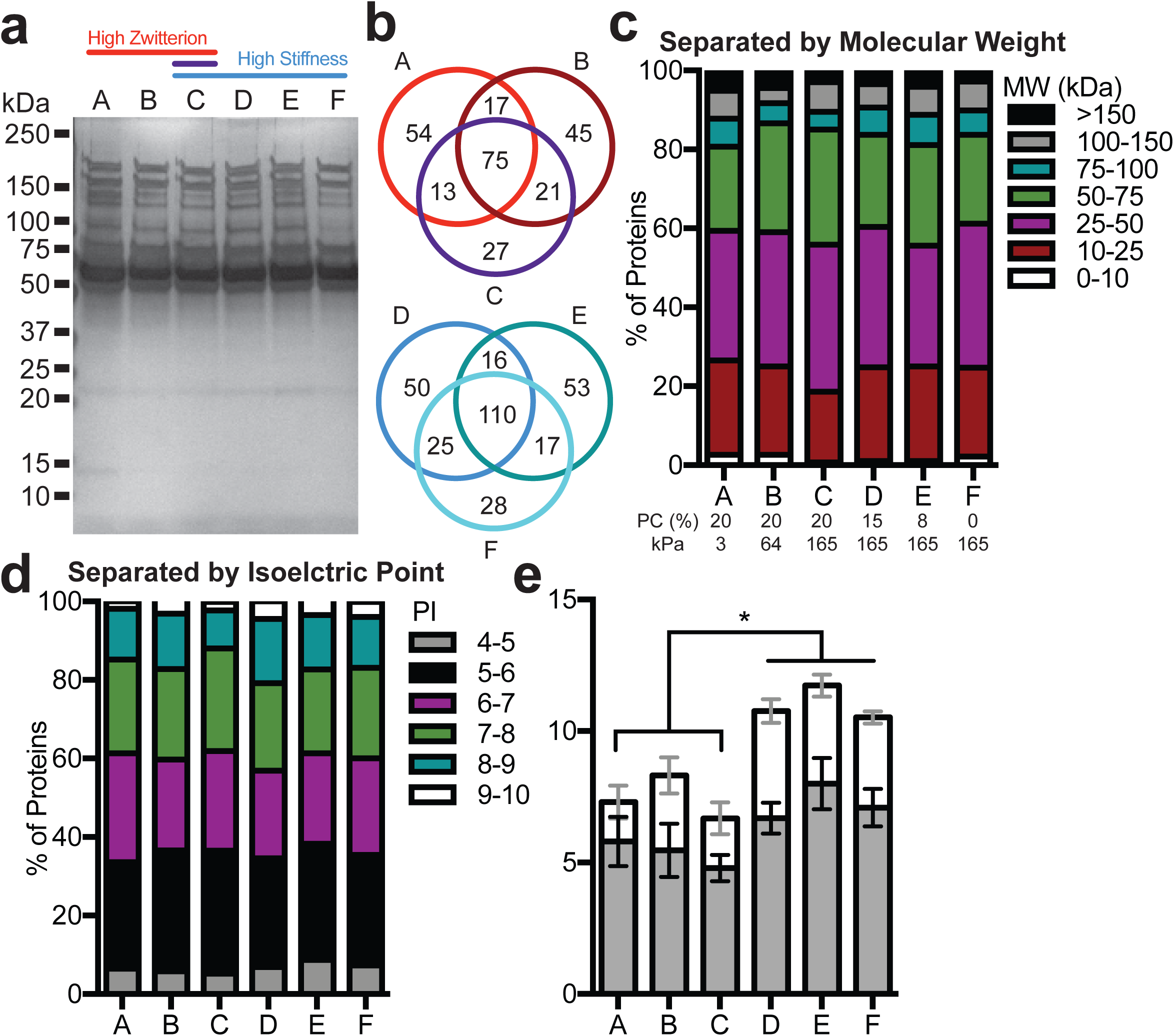
Charged proteins are repelled by highly zwitterionic hydrogels. a) Protein signature was quantified using a silver stain on the *in vivo* proteins adsorbed to each hydrogel. The percentage of PC content and the Young’s modulus for each hydrogel is labeled below. b) Venn diagram depicts the number of shared proteins adsorbed unto the hydrogels. c) Relative percentage of all proteins separated by molecular weight and d) protein isoelectric points, with the e) extreme percentages of isoelectric point data plotted separately. Data is presented as mean with standard deviation (N=3 per hydrogel condition).

**Figure S4.**
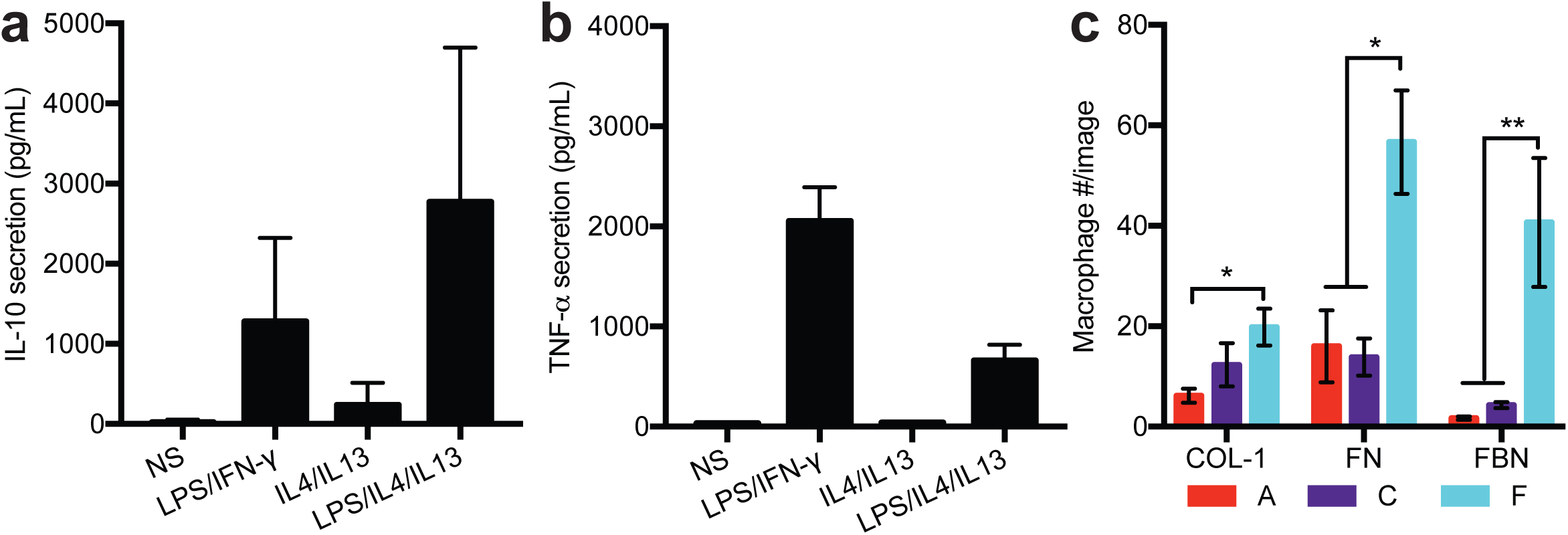
Macrophage response is affected by stimulation on tissue culture plastic. The secretion of a) IL-10 and b) TNF-alpha from macrophages seeded on plastic and stimulated with different combinations of soluble factors. Stimulation of each condition was as follows: NS: no stimulation, LPS/IFN-γ: 1ng/mL LPS and IFN-γ, IL4/IL13: 20ng/mL IL4 and IL13, LPS/IL4/IL13: 0.5ng/mL LPS and 20ng/mL IL4 and IL13. Error bars are the SD (N=2,n=5). c) The number of macrophages adhered after 24 hours to hydrogels exposed to 10 ng/mL of proteins (Col-1: Collagen 1, FN: Fibronectin, FBN: Fibrinogen) for 30 minutes before cell seeding (N=2, n=2). Significance is determined using a two-tailed t-test where p=0.05.

**Table S2.**
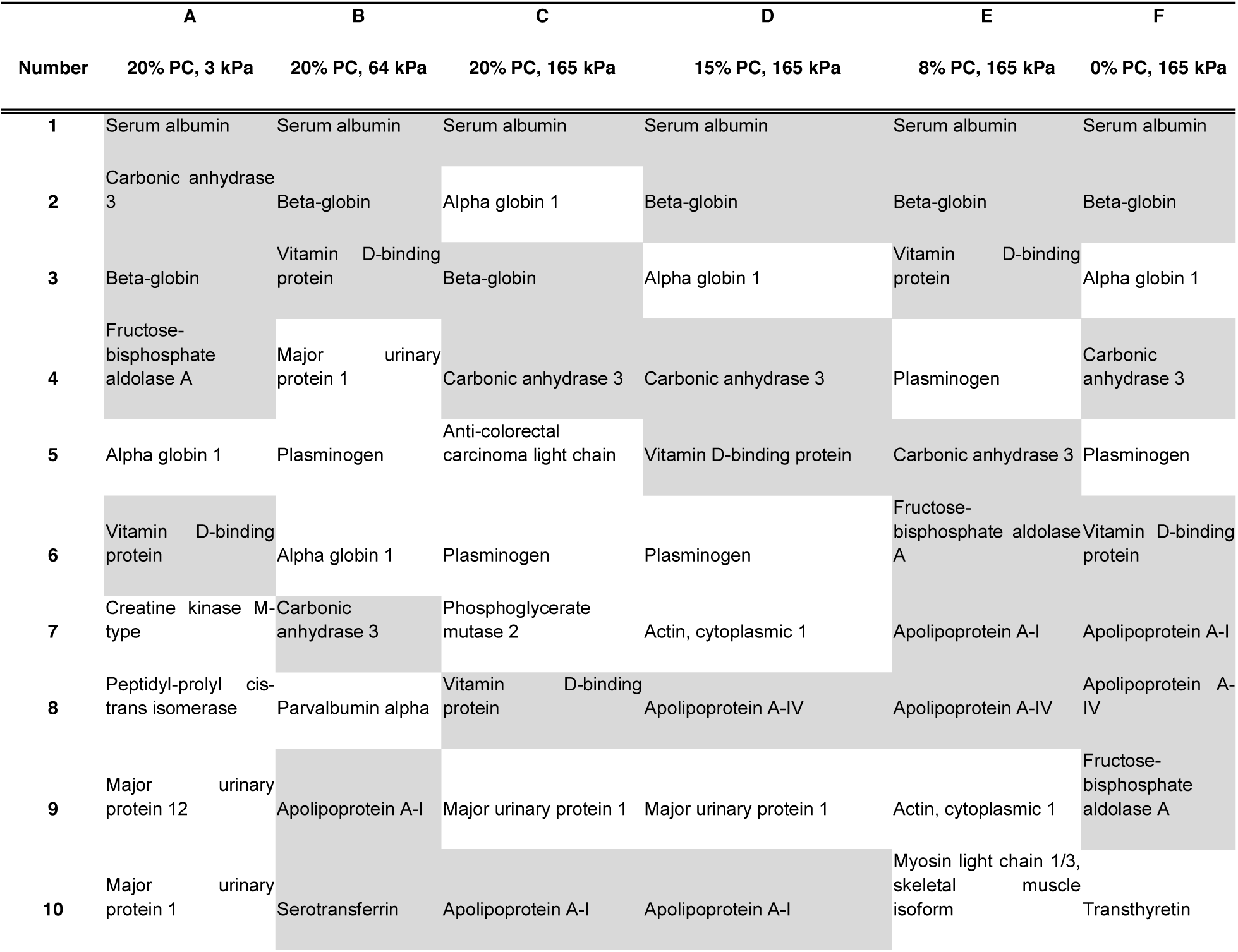
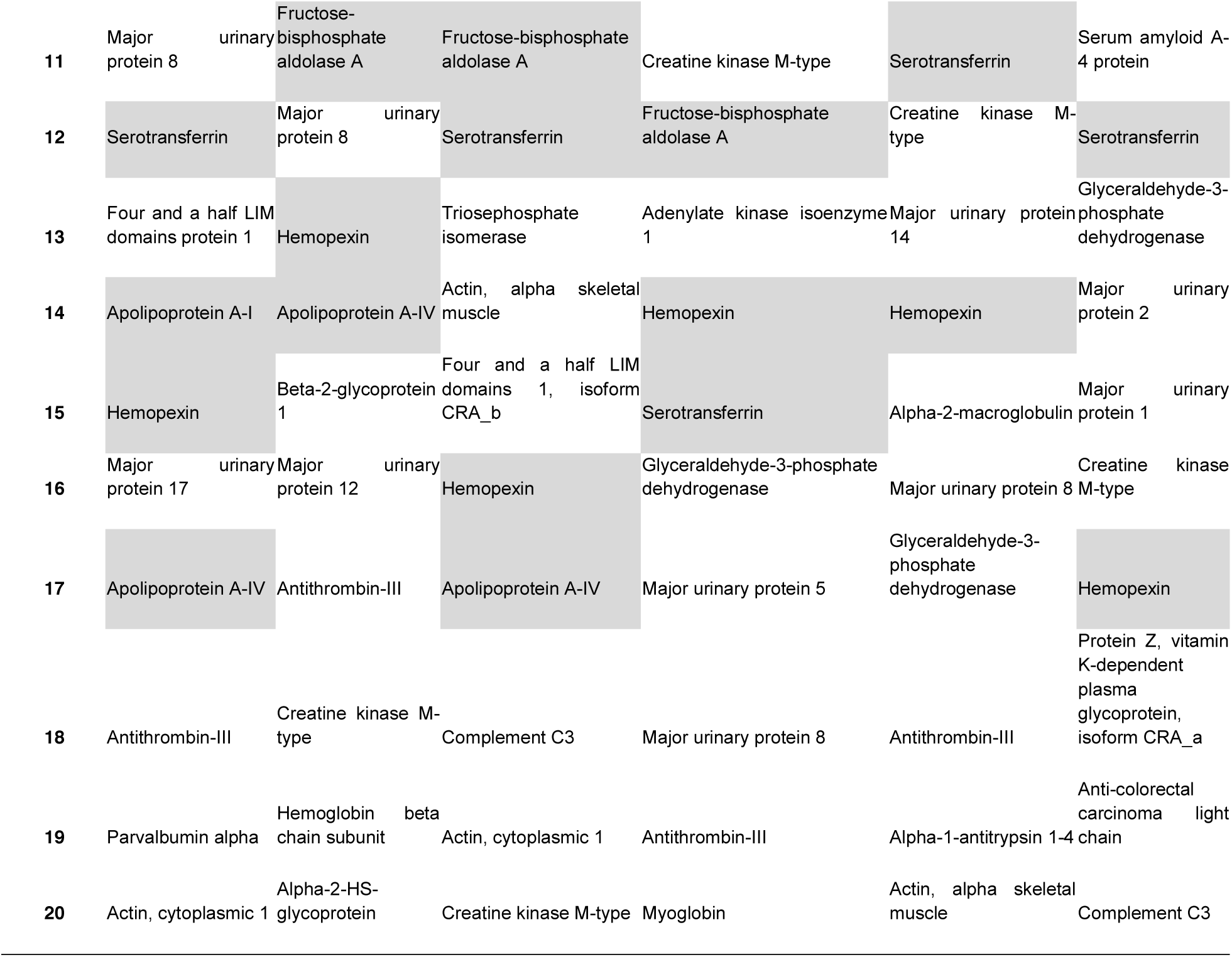
Top 20 protein hits across all platforms. Highlighted proteins were identified in all of the conditions.

